# Extreme Fuzzy Association of an Intrinsically Disordered Protein with Acidic Membranes

**DOI:** 10.1101/2020.09.16.299974

**Authors:** Alan Hicks, Cristian A. Escobar, Timothy A. Cross, Huan-Xiang Zhou

**Author notes:** Equal contribution (Alan Hicks, Cristian A. Escobar). Correspondence should be addressed to H.X.Z.

## Abstract

Many physiological and pathophysiological processes, including *Mycobacterium tuberculosis* (*Mtb*) cell division, may involve fuzzy membrane association by proteins via intrinsically disordered regions. The fuzziness is extreme when the conformation and pose of the bound protein and the composition of the proximal lipids are all highly dynamic. Here we tackled the challenge in characterizing the extreme fuzzy membrane association of the disordered, cytoplasmic N-terminal region (NT) of ChiZ, an *Mtb* divisome protein, by combining solution and solid-state NMR spectroscopy and molecular dynamics simulations. In a typical pose, NT is anchored to acidic membranes by Arg residues in the midsection. Competition for Arg interactions between lipids and acidic residues, all in the first half of NT, makes the second half more prominent in membrane association. This asymmetry is accentuated by membrane tethering of the downstream transmembrane helix. These insights into sequence-interaction relations may serve as a paradigm for understanding fuzzy membrane association.

## Introduction

Upon binding to their partners, intrinsically disordered proteins span a continuum in the extent of structuredness, from fully folded to partially ordered to fully disordered. The “extreme” fuzzy complexes in the latter case have been characterized when the partners are another disordered protein or a nucleic acid^1–5^. A third class of partners for disordered proteins comprise membranes^6–9^. In a well-characterized case, membrane association of α-synuclein is accompanied by the formation of an amphipathic α-helix^9^. A large fraction of transmembrane and peripheral membrane proteins contain disordered regions^10^, but there is little knowledge on any extreme fuzzy complexes with membranes. Here we tackle the challenge of characterizing the extreme fuzzy membrane association of the disordered cytoplasmic N-terminal region of the transmembrane protein ChiZ, a member of the *Mycobacterium tuberculosis* (*Mtb*) divisome complex, by combining solution and solid-state NMR spectroscopy with molecular dynamics (MD) simulations.

Many disordered proteins are enriched in charged residues^11^, and interactions between opposite charged residues are crucial features of extreme fuzzy complexes between disordered proteins^1, 2^. Likewise the interactions between basic residues of proteins and acidic phosphate groups of nucleic acids are crucial for their high-affinity, fuzzy association^3–5^. The inner leaflet of the plasma membrane is highly acidic, due to asymmetric distribution of charged lipids including phosphatidylserine, phosphatidylinositol, and the latter’s phosphorylated variants^12^, and thus forms a target for polybasic proteins including signaling molecules^7^. The *Mtb* inner membrane contains an abundance of acidic lipids, with phosphatidylglycerol, cardiolipin, phosphatidylinositol, and phosphatidylinositol mannosides present at roughly a 7:3 ratio to the neutral phosphatidylethanolamine (based on the composition in *Mycobacterium smegmatis,* a non-pathogenic model ^13^). This acidic surface provides ample opportunities for association by ChiZ and other *Mtb* divisome proteins with disordered cytoplasmic regions that are enriched in basic residues (Fig. 1a and Supplementary Fig. 1).

**Figure 1.**
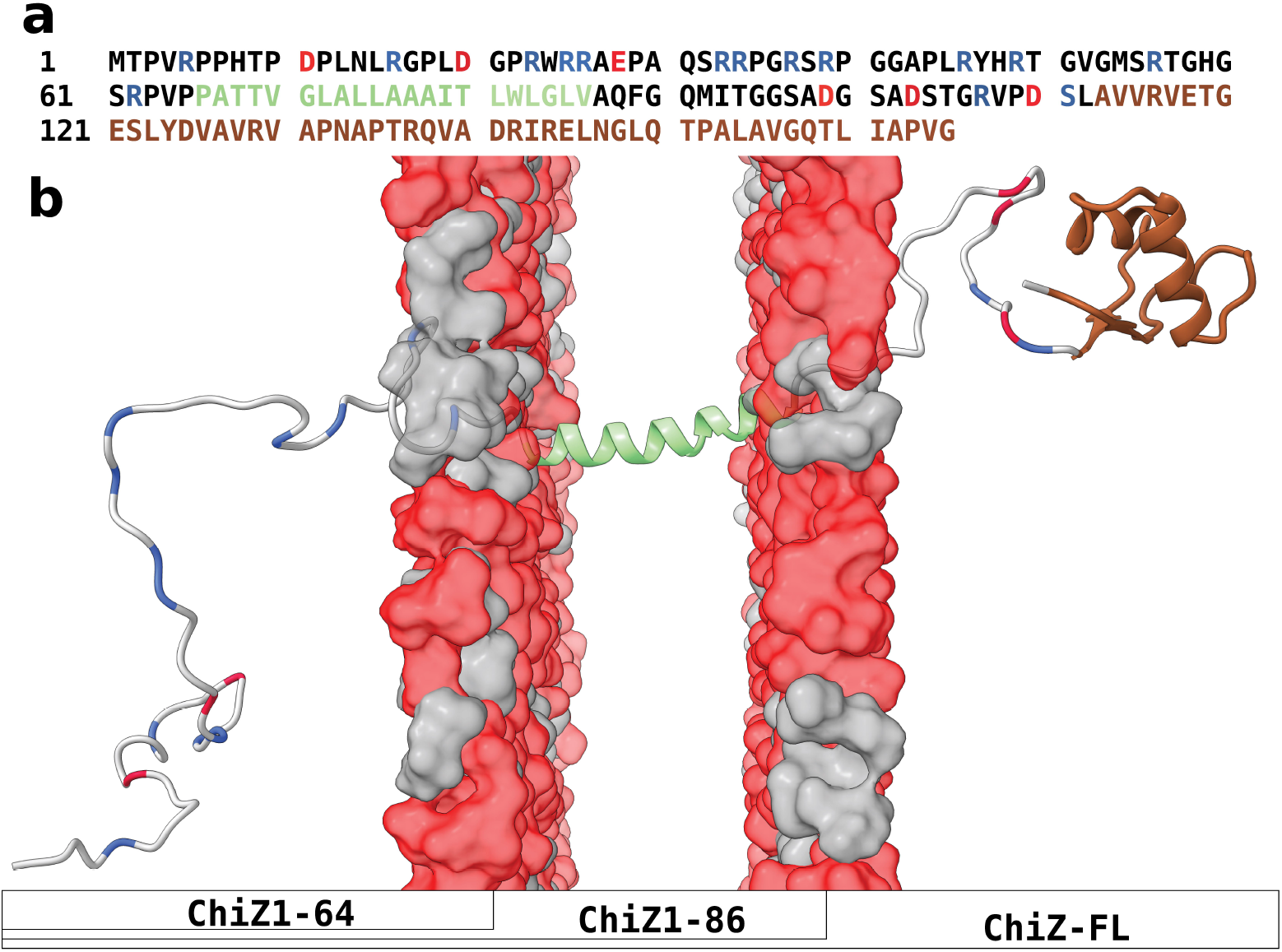
Sequence and structure of ChiZ. (**a**) Amino-acid sequence. Disordered regions, transmembrane helix, and LysM domain are indicated by black, green, and brown letters, respectively; acidic and Arg residues in disordered regions are shown as red and blue letters, respectively. (**b**) Disposition of different regions or domains with respect to the membrane. Colors match those in panel (a). The compositions of the three ChiZ constructs are also indicated.

Very few fuzzy complexes between disordered proteins and membranes have been characterized at the residue level. The most intensely studied protein in this regard is α-synuclein, which forms amphipathic α-helices in the first 100 residues upon membrane association^9^. α-Synuclein preferentially binds to vesicles containing acidic lipids^14^, but membrane curvature also plays an important role. A disease-associated charge reversal, E46K, strengthened membrane binding, but weakened selectivity for membrane curvature^15^. Conversely, increasing negative charges in the C-terminal tail weakened membrane association but enhanced curvature selectivity^16^. The entire 100 residues apparently do not bind to the same vesicle all the time; while the first 30 or so residues stably binds to a vesicle, the remaining segments can dissociate and even bind to a different vesicle, leading to vesicle clustering^17^. Even when the 100 residues were membrane-bound, MD simulations showed significant conformational heterogeneity for α-synuclein, although the helices remained intact^18^. By contrast, no information is available for how a basic region of the Wiscott-Aldritch Syndrome protein interacts with acidic lipids of the plasma membrane, even though the fuzzy interaction activates this protein for stimulating Arp2/3-mediated initiation of actin polymerization^6^. Likewise, the disorder intracellular region of the prolactin receptor, known to interact with inner leaflet-specific lipids via conserved basic clusters and hydrophobic motifs^8^, was modeled without considering membrane association due to lack of information^19^.

Here we report residue-level characterization of the extreme fuzzy association of the ChiZ 64-residue N-terminal region (NT) with acidic membranes. In full-length ChiZ (ChiZ-FL), NT is followed by a 21-residue transmembrane helix; on the periplasmic side, a C-terminal LysM domain (residues 113-165) is connected to the transmembrane helix by a 26-residue linker (Fig. 1). In a previous study ^20^, we showed that, in solution, the NT-only construct, ChiZ1-64, is fully disordered without detectable α-helix or β-sheet formation, but with polyproline II (PPII) formation and intramolecular interactions including salt bridges concentrated in the first half of the sequence. Here we investigated NT-membrane association by solution NMR in the context of ChiZ1-64 and by solidstate NMR on both ChiZ1-64 and ChiZ-FL. In addition, extensive MD simulations of these two constructs and ChiZ1-86 with membranes (Fig. 1b) were carried out, encompassing 16 to 20 trajectories and 20.6 to 38 μs of simulation time for each system.

## Results

### ChiZ1-64 associates with acidic membranes but terminal residues remain free

^1^H-^15^N HSQC spectra of ChiZ1-64 in the presence of liposomes with four lipid compositions were acquired to assess membrane association (Fig. 2). Association is indicated by loss of crosspeaks due to line broadening from the slow tumbling when a residue interacts with a liposome. Relative to the ^1^H-^15^N HSQC spectrum of ChiZ1-64 in solution (hereafter “unbound” ChiZ1-64)^20^, no major loss in NMR signals was detected when the liposomes contained POPC only (Fig. 2a), 4:1 DOPC:DOPE (Fig. 2b), or 4:1 POPC:POPG (Fig. 2c). These spectra show that ChiZ1-64 does not associate significantly with the neutral POPC and DOPC:DOPE membranes, or even with a membrane containing 20% acidic lipids. In contrast, the HSQC spectrum in the presence of 7:3 POPG:POPE liposomes, which mimic the charge composition of *Mtb* membranes^13^, shows that the crosspeaks of most of the residues are broadened beyond detection (Fig. 2d). Of the remaining crosspeaks, based on the overlap with the counterparts in unbound ChiZ1-64, assignments could be made for Gly58, Ser61, Arg62, and Val64, all located at the C-terminus. Due to slight shifts, the few other crosspeaks could not be unambiguously assigned, but appear to be N-terminal residues, including Met1, Thr2, His8, Thr9, and Asn14, as well as possibly Gln31. So the HSQC spectra demonstrate that ChiZ1-64 associates with membranes containing 70% acidic lipids, but residues at the two termini remain free.

**Figure 2.**
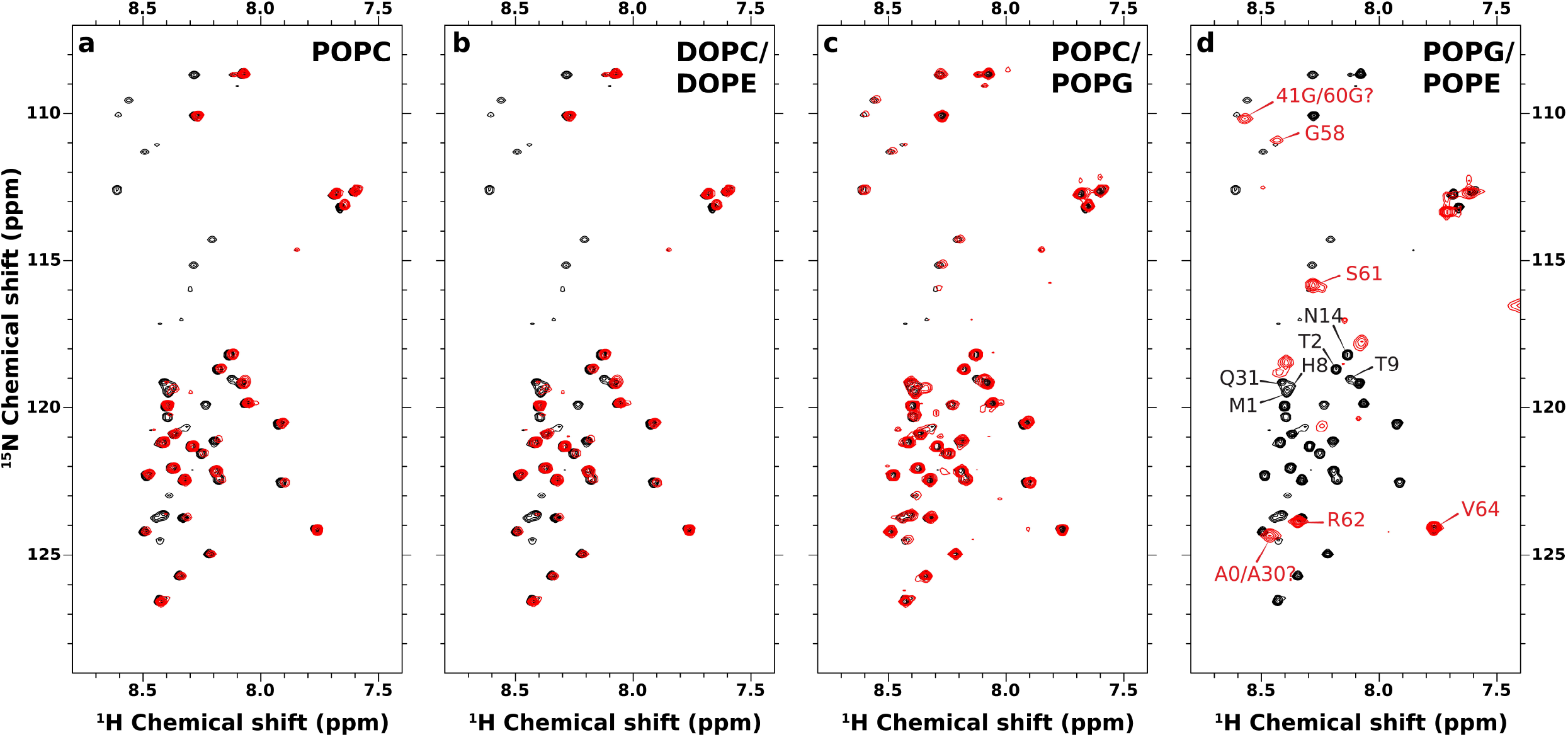
Solution ^15^N-^1^H HSQC spectra of ChiZ1-64 in the presence and absence of liposomes. ChiZ1-64 spectra without and with liposomes are shown as black and red contours, respectively. ChiZ1-64 was mixed with (**a**) POPC, (**b**) DOPC:DOPE (4:1 molar ratio), (**c**) POPC:POPG (4:1 molar ratio), and (**d**) POPG:POPE (7:3 molar ratio) at a protein to lipid ratio of 1:100 in 20 mM phosphate buffer (pH 7.0) containing 25 mM NaCl. All spectra were collected with 100 μM protein at 25 °C.

### Membrane association is fuzzy, but hint for a sub-population with a stable motif

The solution NMR HSQC experiment is useful for indicating membrane association, but the loss of crosspeaks precludes further characterization of the association. We thus turned to magic-angle-spinning (MAS) ^13^C solid-state NMR experiments: insensitive nuclei enhanced by polarization transfer (INEPT) and cross polarization (CP). The former is sensitive to dynamic sites, whereas the latter is sensitive to static sites. The INEPT spectrum of ChiZ1-64 bound to POPG:POPE liposomes shows an abundance of crosspeaks (Fig. 3a, black contours), indicating that most of the residues remain highly dynamic and hence the membrane association is extremely fuzzy. In fact, the dynamics apparently rival those in unbound ChiZ1-64 and result in the same, undispersed chemical shifts for a given pair of carbon-carbon sites (e.g., Arg Cβ-Cδ) at different positions along the amino-acid sequence. This spectral overlap is a strong indication that ChiZ1-64 does not fold upon membrane association, and allowed the assignment of INEPT crosspeaks to types of carbon-carbon sites, but not to specific residue positions.

**Figure 3.**
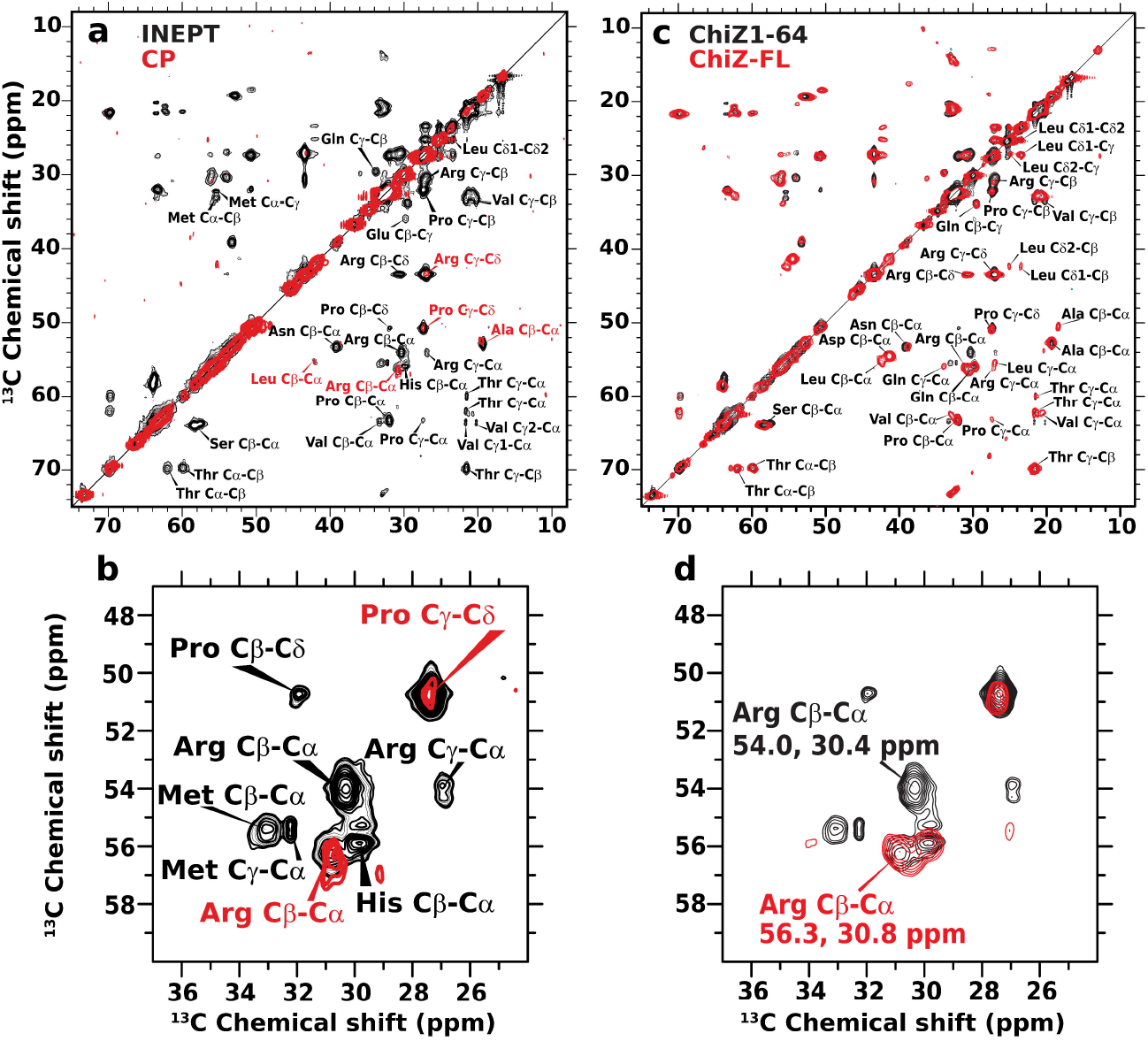
Solid-state NMR data of ChiZ1-64 bound to and ChiZ-FL reconstituted into POPG:POPE liposomes. The protein to lipid ratios were 1:50 and 1:80, respectively, for the two constructs. (**a**) ^13^C-^13^C correlation spectra of ChiZ1-64 using INEPT and CP magnetization transfer, shown in black and red, respectively. The INEPT spectrum was acquired with 512 transients and 128 scans per transient. For the CP spectrum, the PARIS pulse sequence was used with 100 ms mixing time and 400 transients with 268 scans per transient. (**b**) Zoom into a region centered around the Arg Cβ-Cα crosspeaks. (**c**) Comparison of ChiZ1-64 and ChiZ-FL INEPT spectra, shown in black and red, respectively. (**d**) Zoom into the region centered around the Arg Cβ-Cα crosspeaks. All experiments were carried out at 25 °C and at a 12.2 kHz spinning rate.

Both in unbound ChiZ1-64^20^ and in the INEPT spectrum of POPG:POPE-bound ChiZ1-64, the Arg Cα-Cβ pair has two distinct crosspeaks, one at (56.3, 30.8) (chemical shifts in ppm), and the other at (54.0, 30.4) (Fig. 3b, black contours). With the help of MD simulations (Supplementary Fig. 2), we were able to recognize that, in unbound ChiZ1-64, these two crosspeaks were assigned to Arg residues with one distinction: whether the succeeding residue along the sequence is a Pro. We refer these two groups of residues as RP Arg and non-RP Arg, respectively. The (56.3, 30.8) crosspeak belongs to nine non-RP Arg residues whereas the (54.0, 30.4) crosspeak belongs to the four RP Arg residues: Arg5, Arg34, Arg39, and Arg62.

Corroborating the INEPT result that most residues in POPG:POPE-bound ChiZ1-64 are dynamic, the CP spectrum shows only a few crosspeaks (Fig. 3a, red contours). They largely overlap with crosspeaks in the INEPT spectrum, and accordingly can be assigned to Arg Cα-Cβ and Cg-Cδ, Pro Cg-Cδ, Ala Cα-Cβ, and Leu Cα-Cβ. Interestingly, Arg Cα-Cβ appears as a single crosspeak in the CP spectrum (Fig. 3b, red contours); its overlap with the non-RP crosspeak in the INEPT spectrum suggests that this most prominent CP crosspeak comes from one or more non-RP Arg residues. Rather than appearing at isolated positions along the sequence, it is far more likely that the apparently static residues detected by CP form a contiguous stretch for overall stability. There is only a single such stretch, A43PLR46, and the Arg involved is indeed non-RP. The CP experiment thus hints at a stable motif, A43PLR46, that may form in a sub-population of POPG:POPE-bound ChiZ1-64. Our MD simulations sampled a structure for this putative stable binding motif (Supplementary Fig. 3).

We further performed INEPT on ChiZ-FL reconstituted into POPG:POPE liposomes to determine whether NT remained dynamic when the protein was tethered to the membrane via the transmembrane helix. The INEPT spectra of ChiZ1-64 (black contours) and ChiZ-FL (red contours) essentially overlap (Fig. 3c), showing that, in the context of the full-length protein also, NT does not fold upon membrane association. However, one clear distinction emerges for the Arg Cα-Cβ crosspeaks (Fig. 3d). Whereas ChiZ1-64 Arg Cα-Cβ has both an RP crosspeak at (54.0, 30.4) and a non-RP crosspeak at (56.3, 30.8) (red contours), only the latter crosspeak is observed in the ChiZ-FL INEPT spectrum. The disappearance of the RP crosspeak means that the corresponding Arg residues (more precisely, their Cα and Cβ atoms) become more static upon embedding the transmembrane helix in the membrane. Of the four RP Arg residues, rigidification is expected for Arg62, which is right next to the transmembrane helix in ChiZ-FL. The most N-terminal Arg residue, Arg5, could become static because the N-terminal His-tag present in ChiZ-FL (but absent in ChiZ1-64) might attach to the membrane^21^. That still leaves two RP residues, Arg34 and Arg39, in the midsection unaccounted for. As the data from the next experiment indicate, even in the ChiZ1-64 construct, these two residues, along with other midsection Arg residues, interact with lipids and the resulting loss in dynamics might prevent them from being detected by INEPT, but the loss in dynamics was incomplete so Arg34 and Arg39 were not detectable by CP either. Upon embedding the transmembrane helix in the membrane, Arg34 and Arg39 in ChiZ-FL may interact more strongly with lipids and further lose some dynamics (see below), thereby evading detection by INEPT.

### Arg residues engage in direct interactions with lipid headgroups

We used paramagnetic relaxation enhancement to identify NT residues that interact with membranes. By doping liposomes with lipids chelating the paramagnetic ion Gd^3+^ (Fig. 4a), neighboring ChiZ nuclei would relax much faster due to increased dipolar interactions with the spin label, resulted in line broadening and loss of signal intensity. One-dimensional ^13^C direct-excitation spectra show that resonances of Arg side-chain carbons experience significant intensity loss in the presence of Gd^3+^-chelated lipids, while other resonances are largely unaffected (Fig. 4b, c). This observation applies to both ChiZ1-64 bound to POPG:POPE liposomes and ChiZ-FL reconstituted into these liposomes, and reveals that Arg residues are the major players in mediating NT association with membranes.

**Figure 4.**
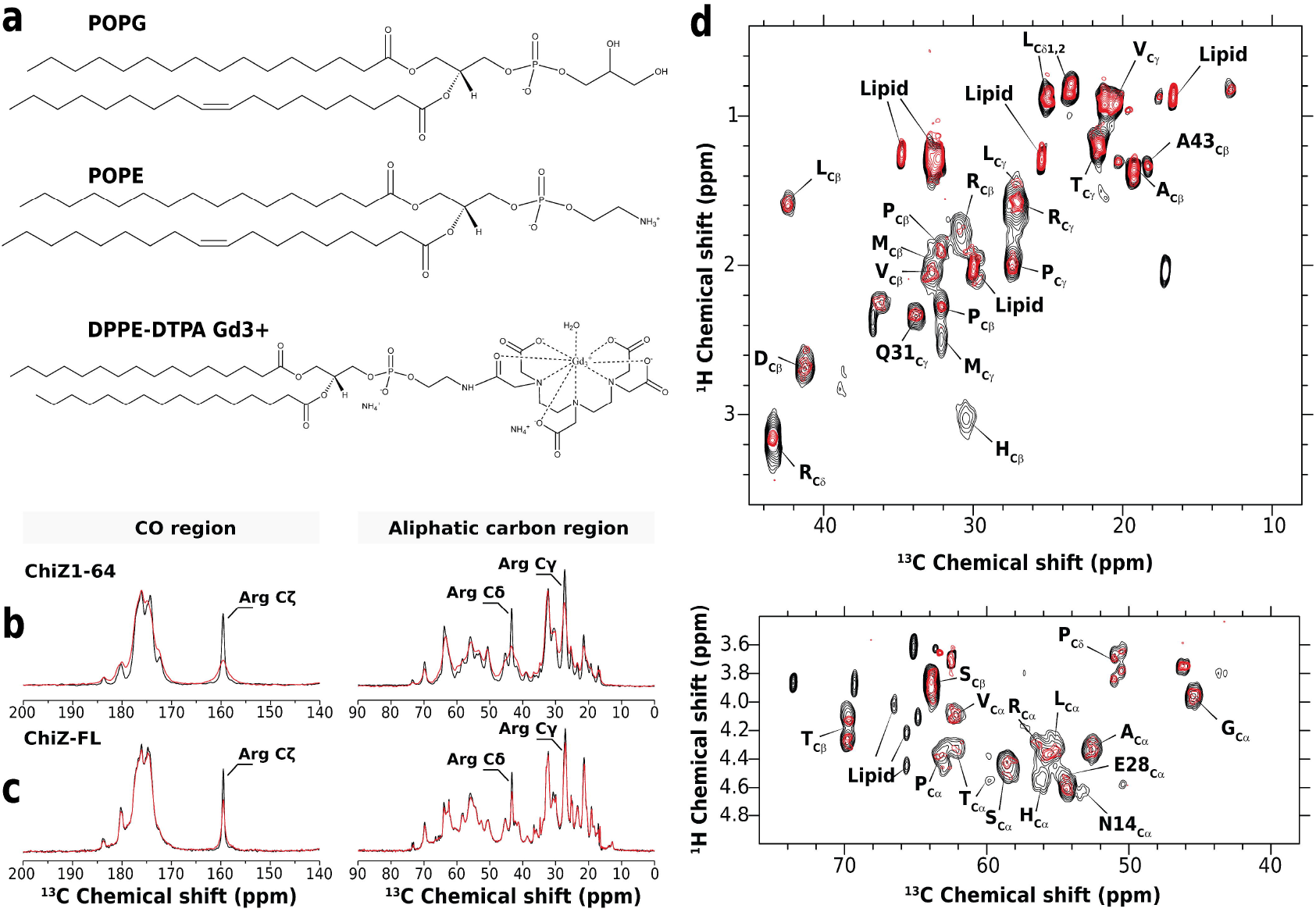
Paramagnetic relaxation enhancement data of ChiZ1-64 bound to and ChiZ-FL reconstituted into POPG:POPE liposomes. The protein to lipid ratios were 1:50 and 1:80, and Gd^3+^-chelated lipids were at 2% and 1%, respectively, for the two constructs. (**a**) Molecular structure of POPG, POPG, and PE-DTPA (Gd). Onedimensional ^13^C direct-excitation spectra of (**b**) ChiZ1-64 and (**c**) ChiZ-FL in the absence (black) and presence (red) of Gd^3+^-chelated lipids. 128 scans were collected on each sample. (**d**) ^1^H-^13^C INEPT-based spectra of ChiZ-FL in the absence (black) and presence (red) of Gd^3+^-chelated lipids. The aliphatic and alpha carbon regions are shown in two panels. All experiments were carried out at 25 °C and at a 12.2 kHz spinning speed.

To characterize NT-lipid interactions in more detail, we investigated paramagnetic relaxation enhancement in reconstituted ChiZ-FL in ^1^H-^13^C correlation experiments with INEPT magnetization transfer. We took advantage of the spectral overlap between the solid-state INEPT and solution HSQC spectra (Supplementary Fig. 4), and assigned the INEPT crosspeaks to types of carbon sites (e.g., Val Cg; Fig. 4d). In a few cases, assignment could be made to specific residues, either because there was only a single residue of a given type (Asn14, Glu28, or Gln31) in NT, or because it was the only NT residue of a given type (Ala43) that preceded a Pro. A comparison of the ^1^H-^13^C correlation spectra between ChiZ-FL samples without (black contours) and with (red contours) the Gd^3+^ spin label provides a global pictures on the NT residues that are in contact with lipid headgroups. An immediate observation is that NT experiences a general loss in ^1^H-^13^C signals in the presence of Gd^3+^. As the relaxation enhancement effect of the spin label may reach protons as far as 20 to 25 Å away, we interpret the general loss in signal as an indication that the spin label senses the entire NT sequence. In other words, when ChiZ-FL is reconstituted into POPG:POPE liposomes, no portion of NT appears to dissociate entirely from membranes.

In the aliphatic region of the ^1^H-^13^C correlation spectra, upon adding the spin label, Arg Cδ sites experience the strongest loss in intensity. In addition, the His Cβ crosspeak disappears altogether. The considerable intensity loss for Arg side chains indicates direct interaction with lipids; the signal disappearance of His side chains likely can be attributed to membrane attachment of the N-terminal His-tag. Similar effects of the spin label on Arg and His residues are also observed in the Cα region of the spectra.

Together, the data from the different NMR experiments indicate that Arg residues away from the NT termini are the major mediators of the association with acidic membranes. The association is extremely fuzzy as NT remains highly dynamic and does not fold, apart from some hint for a sub-population with A_43_PLR_46_ as a stable binding motif.

### NT is anchored to membranes by Arg residues in the midsection

As is clear from the foregoing presentation, our MD simulations were crucial in the interpretation of the NMR data. More importantly, the simulations revealed atomistic details about the extreme fuzzy membrane association of NT, which we now describe. As a first step, we calculated the probabilities that individual NT residues in the three ChiZ constructs are in contact with POPG:POPE membranes (i.e., < 3.5 Å between heavy atoms; Fig. 5a, b). We denote the membrane-contact probability of residue *i* by *C_i_*. In ChiZ1-64, the residues that contact membranes with relatively high probabilities (i.e., *C_i_* > 0.25, indicated by a horizontal dashed line in Fig. 5b) are all Arg residues, in accord with the paramagnetic relaxation enhancement data in Fig. 4a. There are nine such Arg residues, including Arg23, Arg26, Arg33, Arg34, Arg37, Arg39, Arg46, Arg49, and Arg56. In complete agreement with the ^1^H-^15^N HSQC spectra of ChiZ1-64 reported in Fig. 2d, the extreme N- and C-terminal residues do not frequently form contacts with POPG:POPE membranes. Indeed, except for Arg56, the frequent-contact Arg residues are limited to the midsection of NT, with Arg37 having the highest contact probability, at 60%. Furthermore, the distribution of the frequent-contact Arg residues along the sequence gives the first indication that the two halves of NT (denoted as N- and C-half) are not equal in membrane association, with the C-half playing a more prominent role. We will further explore this asymmetry below. A representative snapshot illustrating the membrane anchoring of NT by midsection Arg residues is shown in Fig. 5c.

**Figure 5.**
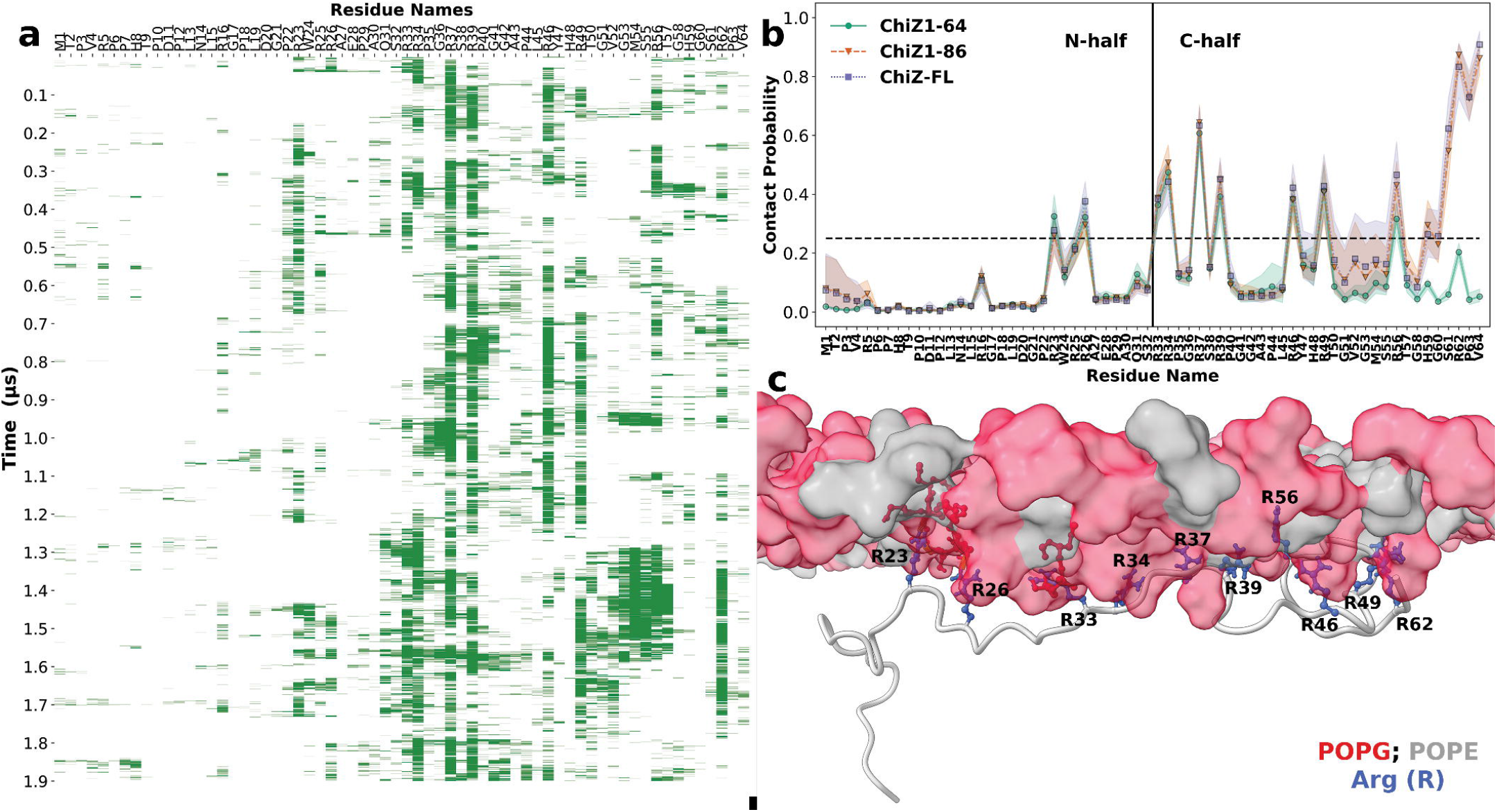
Membrane-contact probabilities of NT residues. (**a**) Contact status of individual residues in snapshots along a 1.9-μs molecular dynamics trajectory of ChiZ1-64. Green bars or blanks indicate that a residue either is or is not in contact with the membrane. (**b**) Membrane-contact probabilities of NT residues in the three constructs. The shaded bands represent standard deviations among the snapshots analyzed. The extreme N-terminal residues that show high membrane-contact probabilities in ChiZ1-86 and ChiZ-FL are from two molecular dynamics trajectories where Met1 was started as nearly embedded in the headgroup region, mimicking in a small way potential membrane attachment of the N-terminal His-tag; Met1 eventually dissociated from the membrane. For these two constructs, residues 49-56 penetrated into the membrane in two trajectories. These events led to relatively large standard deviations in membrane-contact probability. (**c**) A snapshot of ChiZ1-64 at 1.56 μs from the same trajectory as in (a), illustrating the membrane anchoring of NT by Arg residues in the midsection.

Comparing the membrane-contact probabilities of ChiZ1-64 with those of the longer constructs (Fig. 5b), the most obvious effect of membrane tethering of the NT C-terminus is the near 100% contact probabilities of the three most C-terminal residues, R_62_PV_64_. The effect of the membrane tethering is apparent down to residue Thr50, and small increases in membrane-contact probabilities are seen all the way to the start of C-half. These changes accentuate the asymmetry between the two halves of NT in membrane association. Additional evidence below will show that the effect of the membrane tethering even propagates into N-half. The resulting further loss in dynamics for Arg34 and Arg39 in ChiZ-FL explains why they, along with Arg5 and Arg62, are not detectable by INEPT (Fig. 3b).

Lastly, we note that while the membrane-contact probabilities of NT residues are very similar between ChiZ1-86 and ChiZ-FL, there are subtle differences. A majority (11 out of 16) of the frequent-contact residues have slightly higher contact probabilities in ChiZ-FL than in ChiZ1-86 (Supplementary Fig. 5a). This difference will also be further addressed below.

### Competition between acidic residues and POPG is main cause for asymmetry between the two halves of NT in membrane association

The scant involvement in membrane association by Arg residues in ChiZ1-64 N-half stands in contrast to their deep involvement in intramolecular salt bridges when ChiZ1-64 is unbound^20^ (Supplementary Fig. 6). The latter result has been explained by the fact that the salt-bridge partners, i.e., acidic residues (Asp11, Asp20, and Glu28), are all in N-half. Apparently, acidic residues and acidic lipids compete for interactions with Arg residues; when Arg residues (in particular, in N-half) engage in intramolecular interactions with acidic residues, they lose the ability to engage in intermolecular interactions with POPG lipids. Indeed, with the partners being either POPG lipids or acidic residues, the profiles of hydrogen bonding probabilities of Arg residues are mirror images of each other, with N-half favoring acidic residues whereas C-half favoring POPG lipids (Fig. 6a,b).

**Figure 6.**
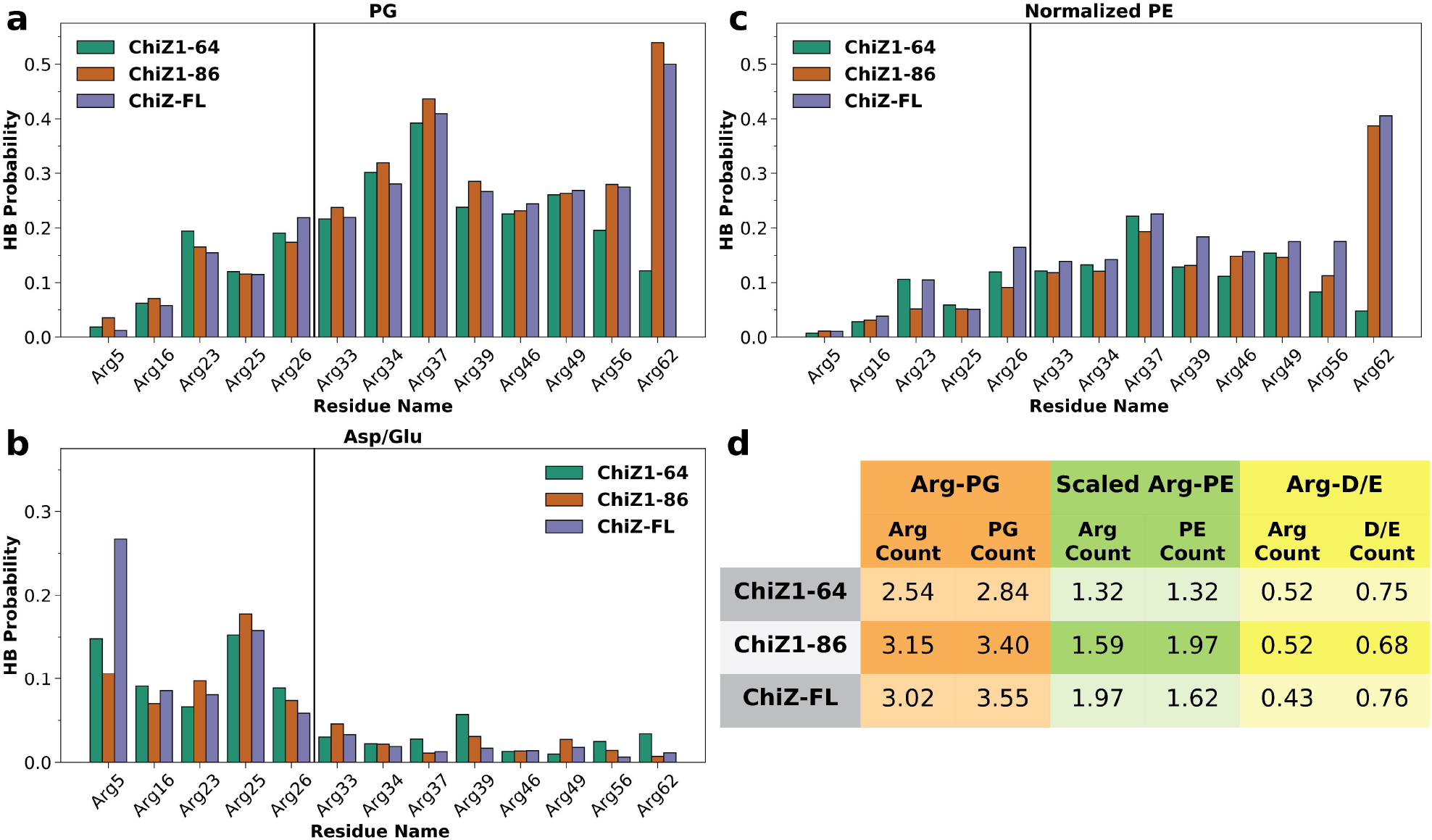
Hydrogen bonding probabilities of NT Arg residues. (**a**) Hydrogen bonding probabilities of Arg residues with POPG lipids. (**b**) Hydrogen bonding probabilities of Arg residues with Asp and Glu residues. (**c**) Hydrogen bonding probabilities of Arg residues with POPE lipids, scaled up by a factor of 7/3. (**d**) Average number of Arg residues that hydrogen bond with a particular type of partner at a given moment, and the counterpart for the partner hydrogen bonding with Arg residues. The partners are either POPG or POPE lipids or Asp and Glu residues.

Expectedly, the probabilities that Arg residues hydrogen bond with POPG lipids (Fig. 6a) track very well their membrane-contact probabilities (Fig. 5b). Indeed, these two sets of data are highly correlated, with a slope of approximately 0.62 (Supplementary Fig. 7). In other words, each time an Arg residue comes into contact with membranes, there is a 2/3 chance that it forms hydrogen bonds with POPG lipids, therefore indicating that Arg-POPG hydrogen bonds as the main driving force for membrane association. Non-Arg residues in ChiZ1-64 have minimal probabilities hydrogen bonding with POPG (Supplementary Fig. 8a). Seven of the nine Arg residues that most frequently hydrogen bond with POPG lipids are in C-half.

In contrast, Arg residues that most frequently hydrogen bond with acidic residues are all in N-half (Fig. 6b). The most prominent of these Arg residues are Arg5 and Arg25. Arg5 can be attributed to its proximity to both Asp11 along the sequence, while Arg25 to its proximity to Asp20 and Glu28. The frequent hydrogen bonding with Asp20 and Glu28 explains why Arg25 has lower probabilities than both of its neighbors, Arg23 and Arg26, for hydrogen bonding with POPG lipids and for membrane contact. Compared to unbound ChiZ1-64 (Supplementary Fig. 6), Arg5 and Arg16 near the N-terminus have increased probabilities of hydrogen bonding with acidic residues upon membrane association, but Arg23, Arg26, and Arg33 have reduced probabilities of hydrogen bonding with acidic residues. Acidic residues thus lose to POPG lipids in their competition for hydrogen bonding with Arg residues.

Besides the acidic POPG, Arg residues can also hydrogen bond with the zwitterionic POPE, though at much lower probabilities (Fig. 6c). Even after compensating for the fact that POPE is at a lower molar fraction of the membranes, Arg residues are still 1.5 to 2.0 times less likely to hydrogen bond with POPE than with POPG (Fig. 6d). On average, 2.5 NT Arg residues in ChiZ1-64 hydrogen bond with POPG lipids at each moment. This number increases to 3.2 in ChiZ1-86 and 3.0 in ChiZ-FL, mostly from C-half Arg residues starting at position 37 (Fig. 6a). In comparison, the average numbers of NT Arg residues that hydrogen bond with POPE lipids at each moment, after scaling up by a factor of 7/3, are only 1.3, 1.6, and 2.0, respectively, in ChiZ1-64, ChiZ1-86, and ChiZ-FL. Therefore POPG lipids preferentially distribute around the membrane-associated NT (see Fig. 5c). Such preferential distribution of acidic lipids around basic groups of membrane-associated proteins have been observed in previous MD simulation studies^22, 23^. On average, each Arg residue engages with 1.1 to 1.2 POPG lipids in their hydrogen bonding. The average numbers of NT Arg residues that hydrogen bond with acidic residues range from 0.52 to 0.43 in the three ChiZ constructs, slightly less than the counterpart, 0.62, in unbound ChiZ1-64.

The two halves of unbound ChiZ1-64 are asymmetric not only in salt-bridge formation but also in PPII propensity (there are very little propensities for helices and β-strands; Supplementary Fig. 9)^20^. Three PPII stretches form with high probabilities (>50%), all in N-half: V_4_RP_6_, P_10_DP_12_, and A_27_EP_29_. In C-half, residues that sample the PPII region with the highest probabilities are P_44_L_45_, at 35%, and S_38_R_39_, at 32%. In agreement with the NMR data, ChiZ1-64 does not gain any secondary structure upon membrane association (Supplementary Fig. 9). In fact, while N-half largely preserves its PPII probabilities upon membrane association, P_44_L_45_ in C-half suffers a modest reduction in its PPII probability, down to 31%. In ChiZ1-86 and ChiZ-FL, this probability further deteriorates to 26% and 25%, respectively. Similar losses in PPII probability are also seen for S_38_R_39_. So NT sacrifices secondary structure in C-half in order to gain stability in membrane association.

The asymmetry in NT’s membrane association is dramatically illustrated by one of the ChiZ1-64 simulation runs (Supplementary Movie 1). In this run, ChiZ1-64 initially binds to one leaflet via N-half. After only 20 ns, it dissociates but then quickly reassociates at 120 ns with another leaflet, this time via C-half. The association is stable for the rest of the 1.9-μs simulation. Apart from the first 20 or residues, NT never dissociates from membranes in the longer constructs.

### Both transmembrane helix and LysM domain contribute, directly or allosterically, to NT-membrane association

Several characteristics of NT-membrane association have emerged from the foregoing analyses of MD simulations. The association is largely maintained by Arg-POPG hydrogen bonding. For ChiZ1-64, these Arg residues are mostly located in the midsection of the sequence, but there is also an asymmetry that favors the C-half. This intrinsic asymmetry is partly due to competition between acidic residues, all in N-half, and POPG lipids for interactions with Arg residues, and partly due to high PPII propensities in N-half. This asymmetry is accentuated by the membrane tethering of the NT C-terminus by the transmembrane helix. As illustrated by Supplementary Movie 2 for ChiZ1-64 and Supplementary Movie 3 for ChiZ-FL, NT-membrane association is highly dynamic. At each given moment, several Arg residues hydrogen bond with the membranes, but the identities of the Arg residues rapidly change (Fig. 5a). As NT changes its conformation and hydrogen bond donors, the lipid acceptors, primarily POPG, also adapt to surround the Arg donors.

To gain a sense of which residues contact membranes at the same time, we calculated the probability, *C_ij_,* that two residues, *i* and *j,* contact membranes at the same time. Fig. 7a-c display the *C_ij_* networks of the three ChiZ constructs as graphs, where circular nodes (with radii proportional to *C_i_*) represent residues with *C_i_* > 0.25, and edge widths represent *C_ij_* (with *C_ij_* threshold at 0.20). It is clear that, relative the contact network of ChiZ1-64, the counterparts of ChiZ1-86 and ChiZ-FL are much more connected, with strong connections extending into N-half. The strengthened network connectivity of the longer constructs arises largely from the higher membrane-contact probabilities of the C-half residues (Fig. 5b), which in turn can be attributed to the membrane tethering of the transmembrane helix. This is the basis of the assertion made above that the effect of membrane tethering propagates all the way into N-half. The direct effect of the membrane-contact probabilities can be removed by normalizing the co-occurrence probability: *Ĉ_ij_* ≡ *C_ij_/C_i_C_j_*, where *C_i_C_j_* is the expected probability that residues *i* and *j* would contact membranes at the same time by chance. A *Ĉ_ij_* that is greater than 1 indicates correlation between the two residues, and hence we refer the *Ĉ_ij_* – 1 network as the contact correlation network. The contact correlation networks no longer show a clear-cut difference in connectivity among nine common residues for ChiZ1-86 and ChiZ-FL and for ChiZ1-64 (Supplementary Fig. 10a-c).

**Figure 7.**
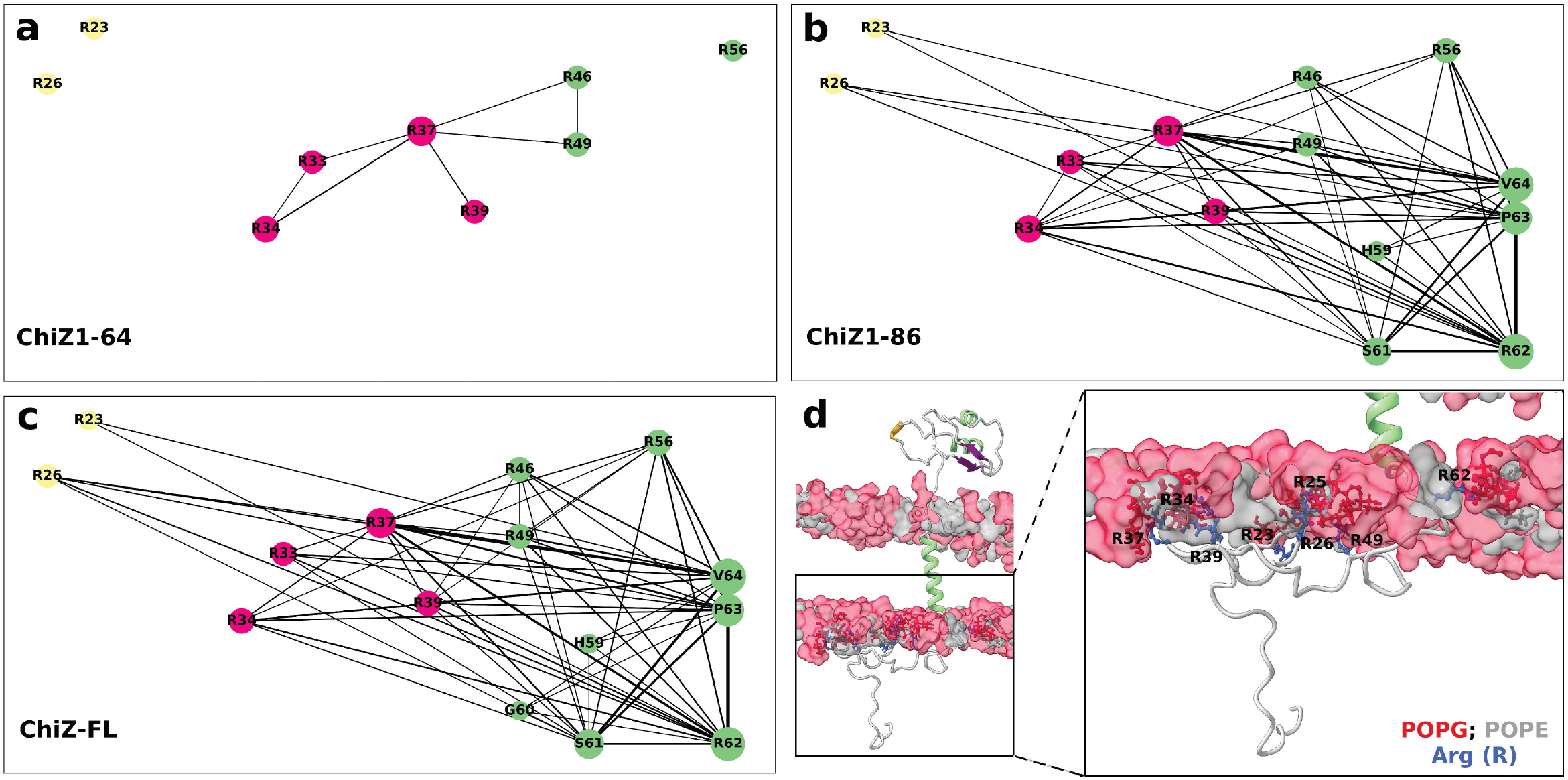
Networks of membrane-contacting residues. (**a-c**) Contact networks of the three ChiZ constructs. Node radii are proportional to contact probabilities *C_i_*; only nodes with *C_i_* > 0.25 are shown. Edge widths are proportional to co-occurrence probabilities *C_ij_*; only edges with *C_ij_* > 0.20 are shown. (**d**) A snapshot of ChiZ-FL illustrating residues that contact the membrane at the same time.

On the other hand, closer inspection reveals that the network connectivity of ChiZ-FL is stronger than that of ChiZ1-86, in line with the consistently higher membrane-contact probabilities of ChiZ-FL shown in Supplementary Fig. 5a. This difference is made clearer by comparing the degree, *d_i_,* defined as the sum of *C_ij_* over all the partner (i.e., *j*) residues, of each node in ChiZ-FL and ChiZ1-86 (Supplementary Fig. 5b). Of the 16 frequent-contact residues, 13 have higher *d_i_* in ChiZ-FL than in ChiZ1-86. As shown by the contact correlation networks (Supplementary Fig. 10b,c), membrane-contact residues in ChiZ-FL also have a higher level of correlation than in ChiZ1-86. The high correlation among NT residues in ChiZ-FL is in accordance with the paramagnetic relaxation enhancement effects observed throughout the NT sequence (Fig. 4d).

The stronger network connectivity of ChiZ-FL reveals that the periplasmic linker and LysM domain also contributes to the stability of NT-membrane association. Periplasmic residues only occasionally contact membranes (Supplementary Fig. 8b), and thus do not influence NT’s membrane association through their own membrane association on the opposite leaflet. Instead, we found that the positioning and tilting of the transmembrane helix are affected by the presence of the periplasmic linker and LysM domain (Supplementary Fig. 11a,b). In ChiZ-FL, the helix shifts toward the periplasmic side by approximately 1 Å, and the helix tilt samples a narrow range of angles. These make the transmembrane helix more deeply (from NT’s perspective) and more stably buried in the membrane. By these changes in the transmembrane helix, the periplasmic linker and LysM domain allosterically strengthen NT-membrane association. Lastly we display a snapshot from the MD simulations of ChiZ-FL in Fig. 7d to illustrate the extreme fuzzy membrane association of NT in the full-length protein.

## Discussion

By combining solution and solid-state NMR spectroscopy and molecular dynamics simulations, we have characterized the extreme fuzzy membrane association of the disordered N-terminal region of ChiZ. The association is largely driven by hydrogen bonding between Arg residues and acidic POPG lipids. Not only the conformation of NT but also the residues that contact the membrane at a given moment are highly dynamic. As NT frolics on the membrane, lipids quickly redistribute, with the acidic POPG lipids preferentially taking up positions next to Arg residues. We refer to membrane association represented by the disordered NT as “semi-specific”, to be contrasted with specific binding between a protein and a macromolecular partner, with a defined interface, and nonspecific binding of proteins at high concentrations, where there is no clear demarcation between a bound state and an unbound state. Membrane association of the disordered NT is also distinct from that of folded domains such as C2 domains in synaptotagmin-1, which have one or more defined membrane-binding sites^23^. For these reasons, ChiZ NT-membrane association represents a new paradigm of bimolecular binding. Other disordered proteins that engage in semi-specific membrane association include α-synuclein^9^ and the Wiscott-Aldritch Syndrome protein^6^.

The term “semi-specific” is also fitting in the sense that NT-membrane association has mixed random and nonrandom characteristics, similar to fuzzy association between two disordered proteins^1, 2^. While the random aspect is obvious from the highly dynamic nature of bound NT (see, e.g., Supplementary Movies 2 and 3), the nonrandom aspect is also worth emphasizing. First, as already noted, it is largely Arg residues that drive the association. Second, for ChiZ1-64, the association-driving Arg residues are located in the midsection of the sequence. Third, the NT sequence codes for asymmetry between the two halves in membrane association. N-half contains all the acidic residues, which compete with POPG lipids for Arg interactions, and has high PPII propensities. N-half is therefore more recalcitrant while C-half is more adaptive to membrane association. Fourth, the intrinsic asymmetry between the two halves of NT is accentuated when its C-terminus is tethered to membranes by the subsequent transmembrane helix. Interestingly, NTs of ChiZ homologues in *Mycobacterium* species have a very conserved C-half, with 6-8 Arg residues (plus a rare Lys residue) and no acidic residues (other than a rare Asp), and a very variable N-half containing all the acidic residues (Supplementary Fig. 12). The characteristics of NT-membrane association determined here for *Mtb* ChiZ thus largely apply to other *Mycobacterium* species, and the conservation of the features important for membrane association argues for a functional role of membrane association.

For specific binding between two structured domains, the dogma is that sequence codes for structure, which in turn codes for specificity, but for fuzzy binding of intrinsically disordered regions including semi-specific membrane association of ChiZ NT and others, how sequence codes for binding specificity is still an open question. Contrary to a-synuclein and other disordered proteins that associate with membranes through amphipathic helices, ChiZ NT does not gain any secondary structure upon membrane association (apart from some hint for an A43PLR46 binding motif in a subpopulation). In the former cases, a mechanism to code for binding specificity is through amino-acid patterning that favors amphipathic-helix formation, i.e., by positive design, as exemplified by the KTKEGV motifs in a-synuclein^9^. In some sense, the specificity of ChiZ NT-membrane association appears to be achieved by negative design. As found in our previous study^20^, the NT sequence codes for correlated segments, mostly in N-half, that are stabilized by salt bridges, cation-π interactions, and high PPII propensities. Just as we speculated previously, these correlated segments lead to the recalcitrance of N-half toward membrane association. Conversely, lack of strongly correlated segments in C-half allows it to be more adaptive to membrane association.

Due to reduced dimensionality, membrane association increases the chances that proteins interact with each other. A main function of ChiZ is to halt cell division, via overexpression, under DNA damage conditions^24^. Overexpression may present ChiZ at a level where NTs of different copies come into contact at membrane. The work presented here characterizing the conformations and dynamics of membrane-bound NT in a single copy of ChiZ lays a solid foundation for understanding interactions between multiple NTs as well as interactions of ChiZ NT and membrane-bound disordered regions of partner proteins including FtsI and FtsQ^25^ (Supplementary Fig. 1).

## Materials and Methods

### Protein expression and purification

Expression and purification of ChiZ1-64 was performed as previously described^20^. ^13^C-^15^N labeled ChiZ-FL containing a non-cleavable N-terminal 6× His-tag was expressed in *Escherichia coli* BL21 Codon Plus RP competent cells. Cells were grown at 37 °C in LB media until OD at 600 nm reached 0.7. Cells were pelleted and transferred to M9 media containing 1 g of ^15^N-ammonium chloride and 2 g of ^13^C uniformly label glucose (Cambridge Isotope Laboratories). After transfer, cells were incubated at 37 °C for 30 min before adding IPTG to a final concentration of 0.4 mM to induce protein expression for 5 hours. Cells were then pelleted and resuspended in a lysis buffer (20 mM Tris-HCl pH 8.0, 500 mM NaCl) for cell lysis using a French press. *n*-Dodecylphosphocholine (DPC; Anatrace) was added to the lysate to a final concentration of 2% (wt/vol) and then incubated overnight at 4 °C with agitation. Cell lysate was centrifuged at 250,000g for 30 min. Protein purification was performed using Ni-NTA resin (Qiagen) equilibrated with the lysis buffer containing 20 mM imidazole. The column was washed using the lysis buffer containing 0.5% (wt/vol) DPC and 60 mM imidazole. Protein was eluted with the same buffer but containing 400 mM imidazole.

### Mixing of ChiZ1-64 with liposomes

ChiZ1-64 was mixed with liposomes containing: (i) pure 1-palmitoyl-2-oleoyl-sn-glycero-3-phosphocholine (POPC); (ii) 1,2-dioleoyl-sn-glycero-3-phosphocholine (DOPC) and 1,2-dioleoyl-sn-glycero-3-phosphoethanolamine (DOPE) at 4:1 molar ratio; (iii) POPC and 1-palmitoyl-2-oleoyl-sn-glycero-3-phospho-(1’-rac-glycerol) (POPG) at 4:1 ratio; or (iv) POPG and 1-palmitoyl-2-oleoyl-sn-glycero-3-phosphoethanolamine (POPE) at 7:3 ratio (all lipids from Avanti Polar Lipids). The protein-liposome mixtures, at a protein to lipid molar ratio of 1:100, were loaded into an NMR tube for collecting ^1^H-^15^N HSQC spectra.

For solid-state NMR experiments, ChiZ1-64 was mixed with POPG:POPE (7:3) liposomes at a 1:50 protein to lipid ratio. The mixture was pelleted down by centrifugation at 15,000g for 15 min. The pellet was then loaded into a 3.2 mm MAS rotor. Liposomes in samples for paramagnetic relaxation enhancement experiments also contained 2% 1,2-dipalmitoyl-sn-glycero-3-phosphoethanolamine-N-diethylenetriaminepentaacetic acid gadolinium salt (PE-DTPA-GD; Avanti Polar Lipids) as a spin label.

### Reconstitution of ChiZ-FL into liposomes

ChiZ-FL samples in MAS solid-state NMR experiments were reconstituted into POPG:POPE (7:3) liposomes at a protein to lipid molar ratio of 1:80. Methyl-β-cyclodextrin (MβCD; Sigma-Aldrich) was used to remove the DPC detergent from the protein-detergent-lipid mixture. Specifically, POPG and POPE lipids in chloroform were mixed and the solvent was removed using nitrogen stream and extensive vacuum. Lipid films were resuspended in 20 mM Tris-HCl (pH 8.0) and sonicated. DPC was added to until the solution became clear. Then ChiZ-FL was added and the mixture incubated for one hour at room temperature. To remove DPC, a solution of MβCD in 20 mM Tris-HCl (pH 8.0) was added to the protein-detergent-lipid mixture at a DPC to MβCD molar ratio of 1:1.5. Proteoliposomes were collected by centrifugation at 250,000g for 3 hours at 8 °C. The pellet was resuspended in 20 mM Tris-HCl (pH 8.0), and an MβCD solution containing 10% of the previous level was added to remove residual detergent. Proteoliposomes were finally collected by centrifugation at 100,000 rpm in a TLA-100 rotor at 8 °C for 16 hours and washed with 20 mM Tris-HCl (pH 8.0) at least twice. ChiZ-FL proteoliposomes were packed into a 3.2 mm MAS rotor for solid-state NMR experiments. Samples for paramagnetic relaxation enhancement experiments were doped with 1% of PE-DTPA-GD.

### NMR spectroscopy

Solution NMR experiments of ChiZ1-64 mixed with liposomes were performed in 20 mM sodium phosphate (pH 7.0) containing 25 mM NaCl, 50 μM sodium trimethylsilylpropanesulfonate (DSS; NMR standard) and 10 % D_2_O. ^1^H-^15^N and ^1^H-^13^C heteronuclear single quantum coherence (HSQC) spectra were collected at 25 °C on an 800 MHz NMR spectrometer equipped with a cryoprobe. Chemical shift assignments of ChiZ1-64 have been reported previously (BMRB accession # 50115)^20^. MAS solid-state NMR experiments of reconstituted ChiZ-FL and liposome-bound ChiZ1-64 were performed at 25°C on a 600 MHz NMR spectrometer equipped with a Low-E MAS probe, with a spinning rate of 12.2 kHz. Glycine carbonyl carbon with a chemical shift frequency of 178.4 ppm was used as ^13^C chemical shift reference. Onedimensional ^13^C direct-excitation spectra were collected using a ^13^C 90° pulse of 62.5 kHz and proton decoupling at 75 kHz using the SPINAL64 decoupling sequence. ^13^C-^13^C (and ^1^H-^13^C) correlation spectra using cross polarization (CP) and INEPT based pulse sequences were collected using the same proton and carbon frequencies as for onedimensional experiments. For CP based experiments, the PARIS pulse sequence was used^26^.

### Molecular dynamics simulations

Three ChiZ constructs (Fig. 1) were modeled and simulated: (i) ChiZ1-64 bound to a 7:3 POPG:POPE bilayer; (ii) ChiZ1-86 with the 22-residue transmembrane helix embedded in a 7:3 POPG:POPE bilayer and NT bound to the inner leaflet; and (3) ChiZ-FL, which extended the ChiZ1-86 system by the periplasmic linker and LysM domain. The simulations of the three systems consisted of 20, 20, and 16 replicate trajectories, respectively; the production lengths of these trajectories were 1.9, 1.8, and 1.29 μs, respectively. The production simulations were preceded by preparatory simulations. The force field combination was AMBER14SB^27^ for proteins, TIP4PD^28^ for solvent (water plus ions), and Lipid17^29^ for membranes.

The membrane-bound ChiZ1-64 simulations were prepared starting from nine ChiZ1-64 models selected from the simulations of the unbound system^20^. A membrane plus solvent system (220 lipids per leaflet with POPG and POPE at 7:3 ratio) was built using the CHARMM-GUI server^30^. The output was converted to AMBER-formatted coordinate and topology files using the *charmmlipid2amber.py* script and tleap in AmberTools17^31^. Upon aligning N-half of ChiZ1-64 to the inner leaflet of the bilayer, ChiZ1-64 was inserted into the system using PARMED, with clashing solvent removed. Using tleap, neutralizing ions plus 25 mM NaCl were added, and the combined system was built into AMBER topology. The final system size was 122 Å × 122 Å × 140 Å with 261,493 atoms.

Preparatory simulations starting from the nine ChiZ1-64 models were run in NAMD 2.12^32^ with AMBER topology. Energy minimization (10,000 cycles of conjugate gradient) was followed by the six-step CHARMM-GUI equilibration protocol^30^, with gradually decreasing restraints on the protein and lipids. Bond lengths involving hydrogens were constrained by the SHAKE algorithm^33^. The timestep was 1 fs in the first four of the six-step protocol but 2 fs in the last two. The durations of the six steps were 25, 25, 25, 200, 200, 2000 ps. Van der Walls interactions were force-switched starting at 10 Å and cut off at 12 Å. The same cutoff was used for calculating short-range electrostatic interactions; long-range electrostatic interactions were treated by the particle mesh Ewald method^34^. The first three steps were under constant temperature (300 K) and volume whereas the last three were under constant temperature and pressure (1.0 atm). Temperature was regulated by the Langevin thermostat with a friction coefficient of 1.0 ps^-1^; pressure was regulated by the Langevin piston^35^ with an oscillation period of 50.0 fs and decay of 25.0 fs. Here and below, whenever pressure was regulated, semi-isotropic scaling in the *x-y* plane was applied to maintain the constant ratio of the two dimensions, with no added surface tension. Following the six-step equilibration, the nine simulations continued under constant temperature and pressure for 40 ns. A total of 20 snapshots, i.e., the nine at the start and the nine at the end of 40-ns simulations, plus two in between, were restarted to run AMBER production simulations for 1.9 μs on GPUs (see below for further details).

ChiZ1-86 models were built using MODELLER^36^, with residues 65-86 modeled as a helix. 10 models were selected for insertion into a POPG:POPE bilayer (at 7:3 ratio with a total of 220 lipids per leaflet) using CHARMM-GUI, with 25 mM NaCl and neutralizing ions added. The final system size was 135 Å × 135 Å × 251 Å with 446,510 atoms. Preparatory simulations of the 10 ChiZ1-86 models were the same as for ChiZ1-64, with the following exceptions. (i) Pressure was regulated by the Monte Carlo barostat; (ii) the durations of the last three steps of the equilibration were 100 ps each; (iii) The subsequent NAMD run was replaced by an AMBER GPU simulation of 1 ns. The 10 final snapshots were each restarted with two random seeds to run AMBER production simulations for 1.8 μs on GPUs.

ChiZ-FL models were built using MODELLER, by combining 8 homology models of the LysM domain (residues 113-165) from SWISS-MODEL^37^ with eight of the ChiZ1-86 starting models. The rest of the ChiZ-FL preparations was the same as for ChiZ1-86. The system contained 300 lipids per leaflet (with POPG and POPE at 7:3 ratio), with a size of 147 Å × 149 Å × 235 Å and 522, 825 atoms. Each of the eight final snapshots in the preparatory simulations was restarted with two random seeds to run AMBER production simulations for 1.29 μs on GPUs.

Production simulations were on GPUs using *pmemd.cuda*^38^ in AMBER18. Temperature was held at 300 K using the Langevin thermostat with a friction coefficient at 1.0 ps^-1^. Pressure was held at 1.0 atm using the Berendsen barostat^39^. For van der Waals interactions, the force-switch distance was 9 Å and cutoff was 11 Å. The latter was also used for dividing direct calculation of electrostatic interactions from a particle mesh Ewald treatment. Bond lengths involving hydrogens were constrained by the SHAKE algorithm. The timestep was 2 fs. Snapshots were saved every 10 ps in the ChiZ1-64 simulations and every 20 ps in the ChiZ1-86 and ChiZ-FL simulations. The first 2000 saved snapshots for each system were discarded.

### MD trajectory analyses

Heavy atom contacts, hydrogen bonds, distances along the *z* axis, secondary structures were calculated with *cpptraj*^40^. Further analyses and plotting were performed using in-house python scripts. Two heavy atoms between a protein and lipids were in contact if they were within 3.5 Å. Hydrogen bonds were defined as formed when the donor-acceptor distance was less than 3.5 Å and the donor-hydrogen-acceptor angle was greater than 135°.

The membrane-contact probability *C_i_* and the probability *C_ij_* that two residues contact membranes at the same time were calculated after pooling all the saved snapshots of each system. From *C_i_*, *C_ij_,* and *C_ij_/C_i_C_j_* – 1, the python module *networkx* was used to build the membrane contact networks and the contact correlation networks. The SHIFTX2^41^ software was used to calculate the chemical shifts of all atoms on snapshots taken at 200-ps intervals. The *seaborn* plotting module in python3 was implemented to create the violin plots. Images of structures were rendered using ChimeraX^42^ and movies were composed using Blender^43^.

## Acknowledgments

A.H. would like to thank Dr. Sean Smrt and Dr. Rongfu Zhang for discussions on NMR. This work was supported by National Institutes of Health Grants R35 GM118091 and R01 AI119178. The NMR experiments were performed at the National High Magnetic Field Laboratory, funded by the National Science Foundation Division of Materials Research (DMR-1644779) and the State of Florida.

## Author contributions

A.H., C.A.E., T.A.C., and H.-X.Z. designed the research. A.H. and C.A.E. performed the research and analyzed the data. A.H. and H.-X.Z. wrote the manuscript.

