## Supplementary Figures 1-12 for "Extreme Fuzzy Association of an Intrinsically Disordered Protein with Acidic Membranes"

*Supporting Information*

CrgA (P9WP57)
MPKSKVRKKN DFTVSAVSRT PMKVKVGPSS VWFVSLFIGL MLIGLIWLMV
FQLAAIGSQA PTALNWMAQL GPWNYAIAFA FMITGLLLTM RWH
(+8e;-1e)

FtsI (LOT911)
MSRAAPRRAS QSQSTRPARG LRRPPGAQEV GQRKRPGKTQ KARQAQEATK
SRPATRSDVA PAGRSTRARR TRQVVDVGTR GASFVFRHRT GNAVILVLML
VAATQLFFLQ VSHAAGLRAQ ...
(+24e;-4e)

FtsK (P9WNA3)
MSSKTVARSG TRTSRSKATS RGASRSARSA VPRKRSRPVK GVGRPSRRHH
RSLLVSTGLA CGRAMRAVWM MAAKGTGGAA RSIGRARDIE PGHRRDGIAL
VLLGLAVVVA ASSWFDAA.. ...
(+25e;-3e)

FtsQ (P9WNA1)
MTEHNEDPQI ERVADDAADE EAVTEPLATE SKDEPAEHPE FEGPRRRARR
ERAERRAAQA RATAIEQARR AAKRRARGQI VSEQNPAKPA ARGVVRGLKA
LLATVVLAVV GIGLGLALYF TPAMSAREIV ...
(+21e;-20e)

FtsW (P9WN97)
... TGLQLPLISA GGTSTAATLS LIGIIANAAR HEPEAVAALR
AGRDDKVNRL LRLPLPEPYL PPRLEAFRDR KRANPQPAQT QPARKTPRTA
PGQPARQMGL PPRPGSPRTA DPPVRRSVHH GAGQRYAGQR RTRRVRALEG
QRYG
(+26e;-9e)

black: cytoplasmic; underline:predicted disorder;
green: predicted TM; brown: periplasmic

**Supplementary Figure 1. Enrichment of basic residues in the disordered cytoplasmic regions of five *Mycobacterium tuberculosis* divisome proteins (related to Fig. 1).** Sequences were from Uniprot, with accession numbers given in the parentheses. Disorder was predicted by PONDR-VSL2[1], with the threshold set at 0.50. Transmembrane regions were predicted using the TMHMM v. 2.0[2]. In predicted disordered cytoplasmic regions, basic residues are colored blue and acidic residues are colored red; the total numbers of basic and acidic residues are summarized in the form (+*n*e; –*m*e). The disordered N-terminal region of FtsQ shows a clear separation of acidic and basic residues, concentrating in the first and second halves, respectively.

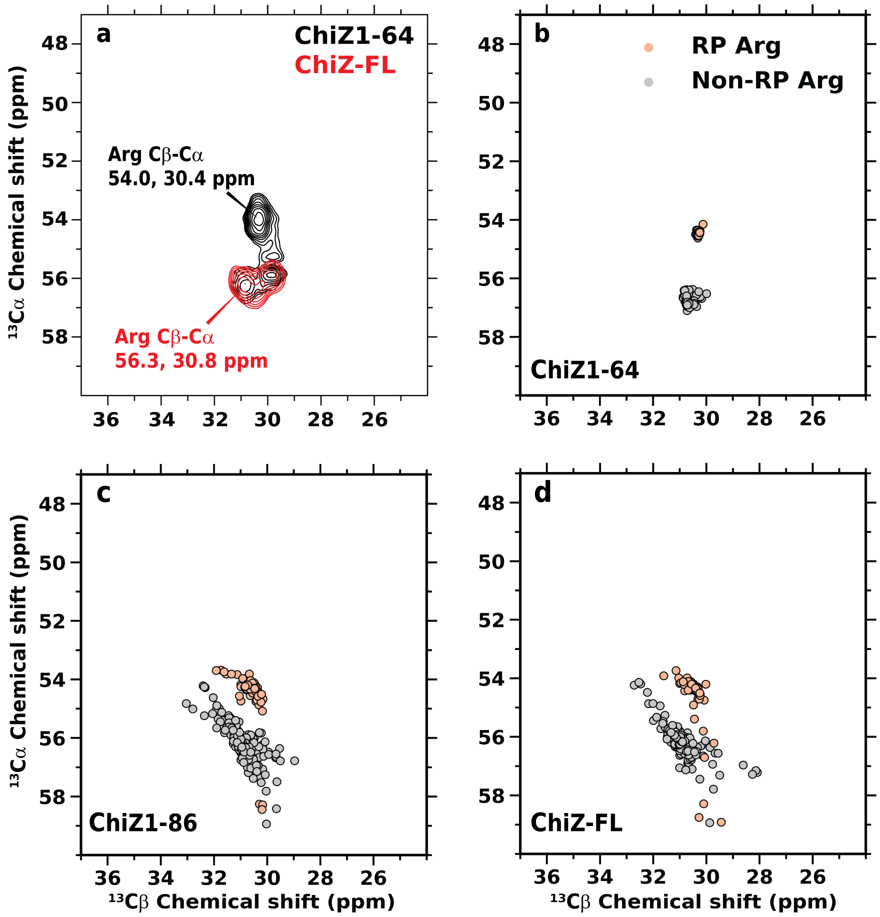


**Supplementary Figure 2. Two distinct sets of Arg (Cβ, Cα) chemical shifts (related to Fig. 3).** (a) $^{13}C$-$^{13}C$ correlation spectra of ChiZ1-64 (black contours) and ChiZ-FL (red contours) using INEPT magnetization transfer, showing only the Arg Cβ-Cα crosspeaks. (b) Predicted Arg (Cβ, Cα) chemical shifts from molecular dynamics simulations of membrane-bound ChiZ1-64, colored pink and gray, respectively, according to whether the Arg residue precedes a Pro residue along the sequence (RP or non-RP). The four RP residues are Arg5, Arg34, Arg39, and Arg62. The average for each Arg residue in each of 20 replicate trajectories is displayed as a circle. The RP cluster corresponds to the (54.0, 30.4) ppm crosspeak in panel (a), whereas the non-RP cluster corresponds to the (56.3, 30.8) ppm crosspeak. (c-d) Corresponding results for ChiZ1-86 and ChiZ-FL. In a few cases, Arg62 moved away from the RP cluster into the non-RP cluster, as a result of the membrane tethering by the subsequent transmembrane helix.

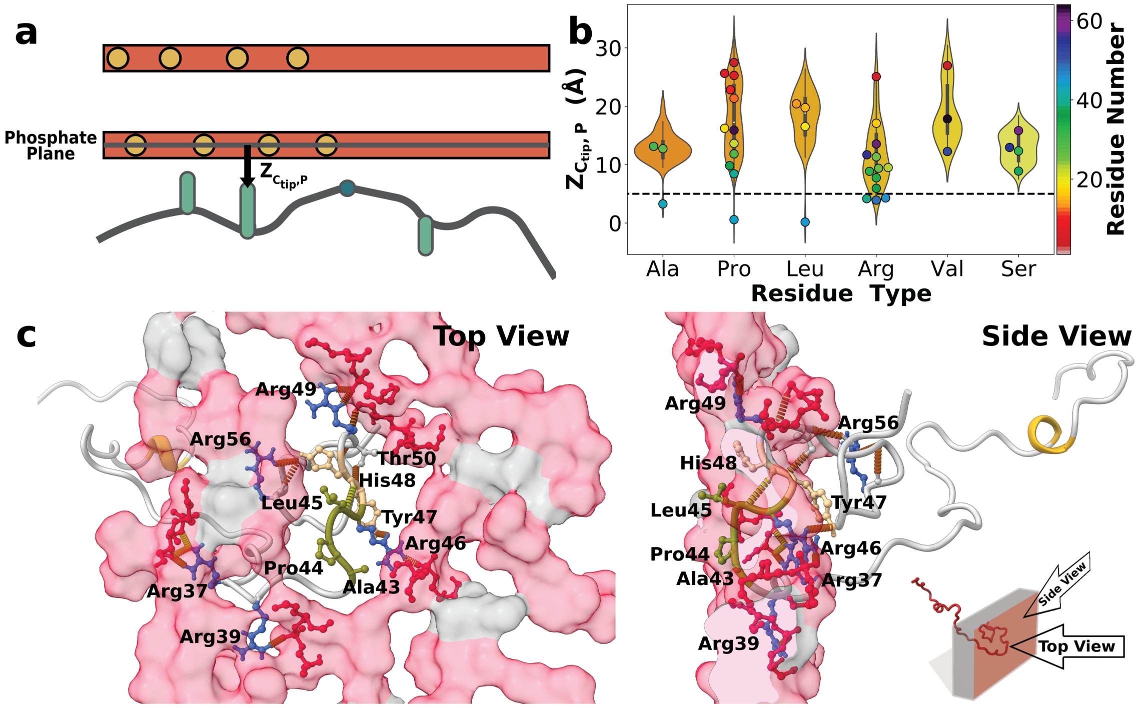


**Supplementary Figure 3. A sub-population with a stable $A_{43}PLR_{46}$ binding motif in ChiZ1-64 (related to Fig. 3).** (a) Illustration of $Z_{C_{tip},P}$, the *z* distance of a side-chain tip carbon from the phosphate plane in the inner leaflet. (b) Statistics of $Z_{C_{tip},P}$ presented as violin plots. A violin plot has two distinct features. In the middle is a boxplot showing the mean of a given residue type (e.g., Ala) over all the sequence positions (e.g., at 27, 30, and 43 for Ala) and over all the simulations, the upper and lower quartiles, and outliers. The wing on each side displays the histogram of $Z_{C_{tip},P}$. Results are shown for the four residue types, Ala, Pro, Leu, and Arg, that were detected in the CP spectrum as well as two other residue types, Val and Ser, for contrast. Three of the CP-detected residue types have outliers below 5 Å (indicated by a dotted line), while the fourth, Arg, samples $Z_{C_{tip},P}$ < 5 Å with a significant probability. The outliers are all from a single trajectory. The mean $Z_{C_{tip},P}$ value for each residue (e.g., Ala27) in that trajectory is shown as a circle, filled by a color according to the position along the sequence (see color scheme on the right). The outlying Ala, Pro, and Leu residues are contiguous in sequence, from Ala43 to Leu45; the next residue, Arg46, also has a mean $Z_{\mathrm{C_{tip}},\mathrm{P}}$ < 5 Å in the same trajectory. $A_{43}PLR_{46}$ thus forms a stable membrane-binding motif in this trajectory. (c) Structure of the $A_{43}PLR_{46}$ binding motif. $A_{43}PL_{45}$ is deeply inserted into the membrane. After sneaking out to partially expose Arg46, which interacts with lipid headgroups, the chain returns with His48 forming a backbone hydrogen bond with Leu45. Additionally, Arg46 Nε hydrogen bonds with Tyr47 Oη.

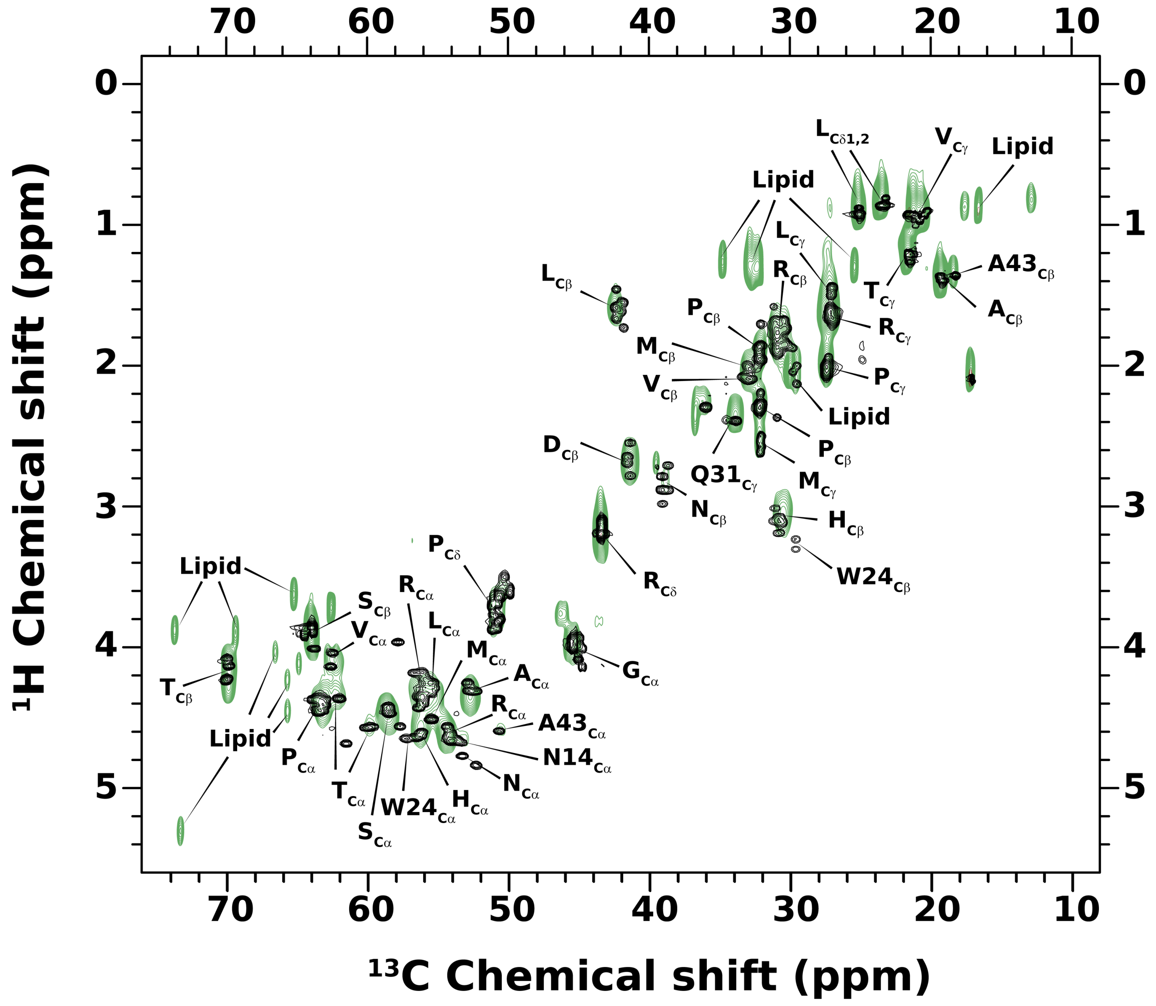


**Supplementary Figure 4. Overlay of $^{1}$H-$^{13}$C correlation spectra of ChiZ1-64 in solution and ChiZ-FL reconstituted into POPG:POPE liposomes (related to Fig. 4).** The solution HSQC spectrum of ChiZ1-64 is shown in black while the INEPT spectrum of ChiZ (at 1:80 protein to lipid ratio) is shown in green.

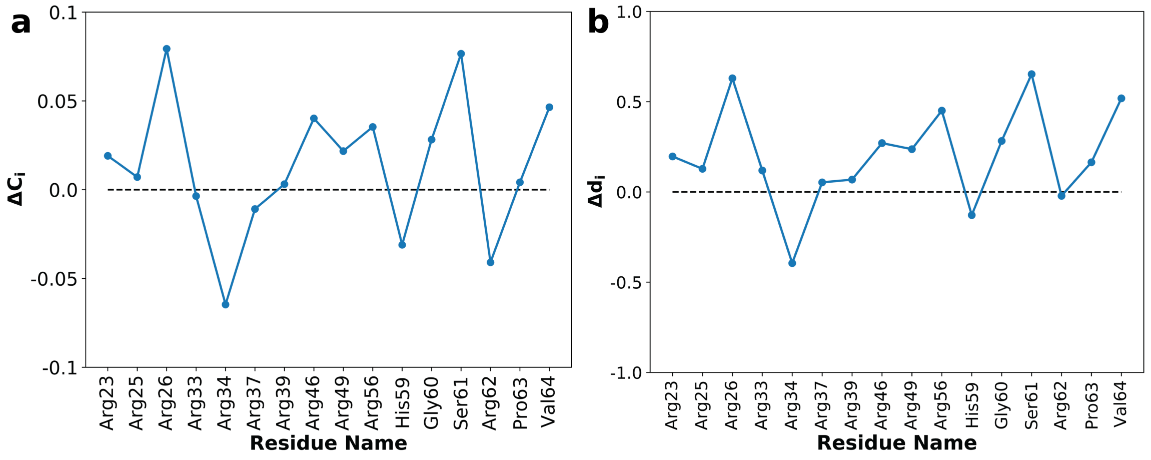


**Supplementary Figure 5. Differences in membrane-contact probability and in degree of network connectivity between ChiZ-FL and ChiZ1-86 (related to Figs. 5 and 7).** (a) Difference in membrane-contact probability, $C_i$, between ChiZ-FL and ChiZ1-86. (b) Difference in degree, $d_i$, between ChiZ-FL and ChiZ1-86.

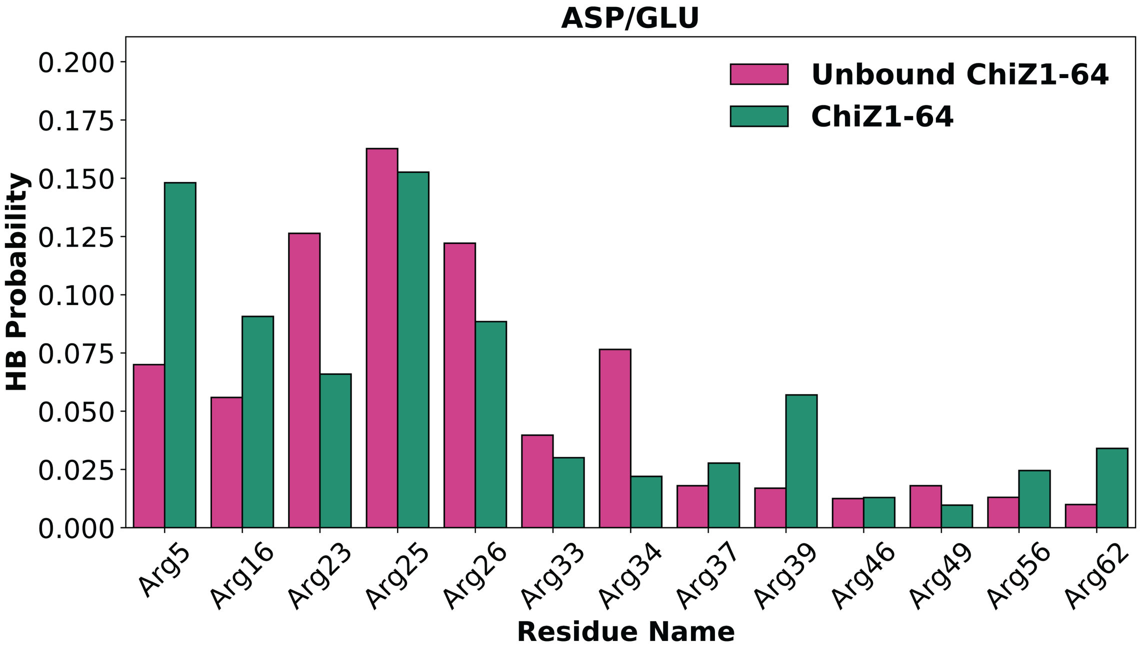


**Supplementary Figure 6. Hydrogen-bonding probabilities of Arg residues with acidic residues in ChiZ1-64, either unbound or membrane-bound (related to Fig. 6).**

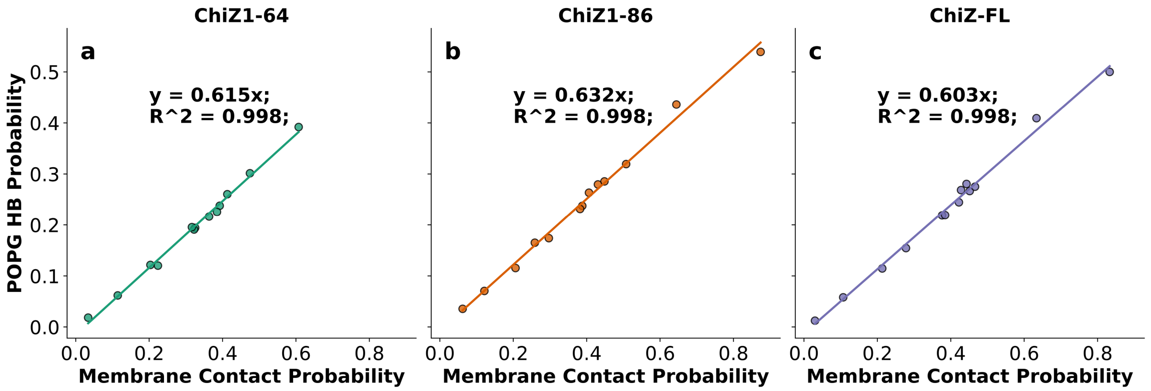


**Supplementary Figure 7. Correlation between membrane-contact probability and hydrogen-bonding probability (related to Figs. 5 and 6).** Each circle represents one of the 13 Arg residues in NT; the hydrogen bonding partner is POPG. (a) ChiZ1-64. (b) ChiZ1-86. (c) ChiZ-FL.

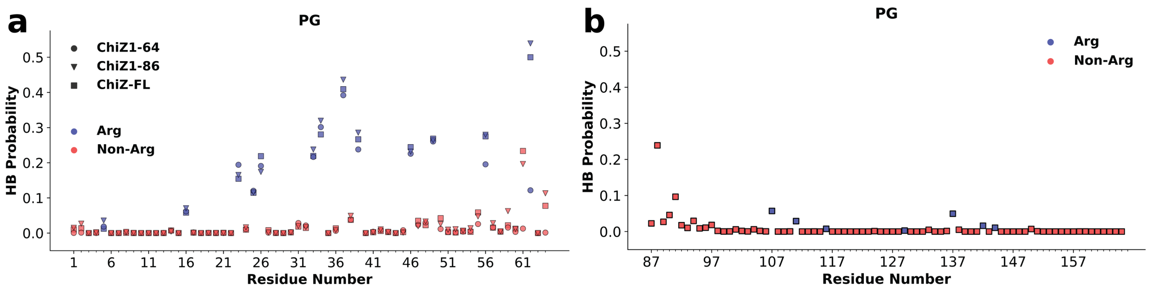


**Supplementary Figure 8. Hydrogen-bonding probabilities of all residues with POPG (related to Fig. 6).** (a) NT residues in the three ChiZ constructs. (b) Periplasmic residues in ChiZ-FL.

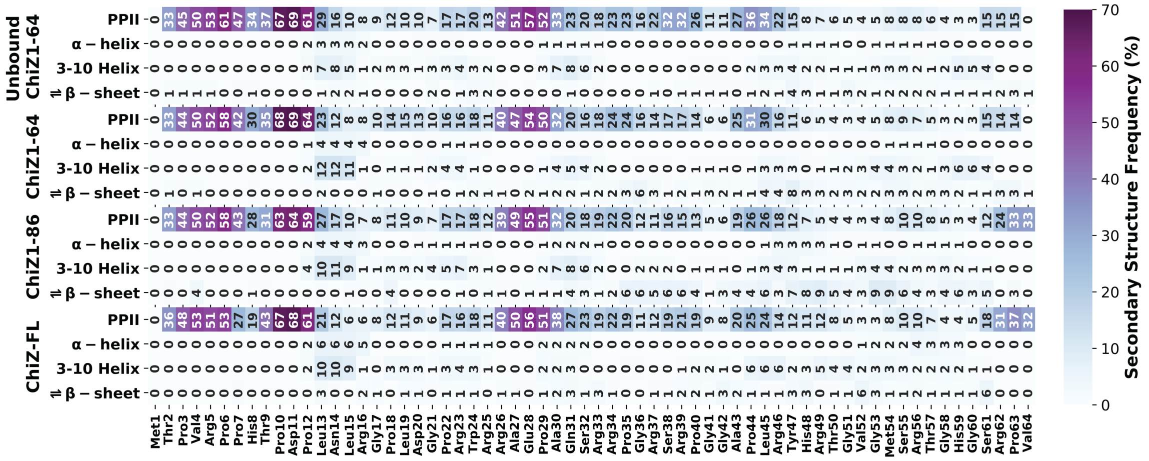

**Supplementary Figure 9. NT secondary structures in three membrane-bound constructs and in unbound ChiZ1-64 (related to Fig. 6).**

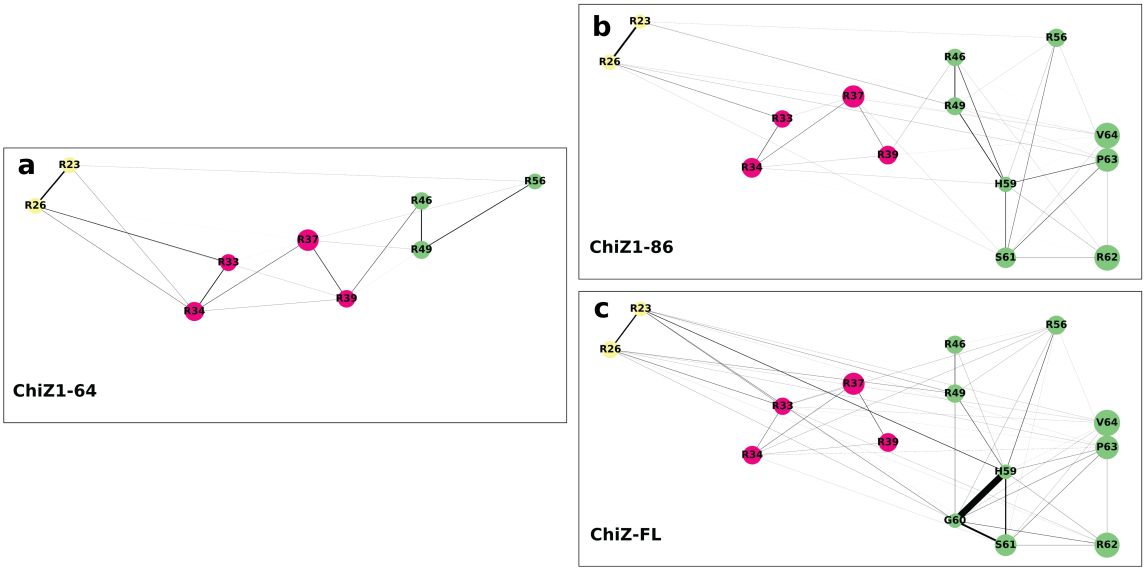


**Supplementary Figure 10. Contact correlation networks of three ChiZ constructs (related to Fig. 7).** Residues with $C_i$ greater than 0.25 are shown as circles (with radii proportional to $C_i$) and edges with $\hat{C}_{ij}$ values with greater than 1 are shown (with width proportional to $\hat{C}_{ij} - 1$).

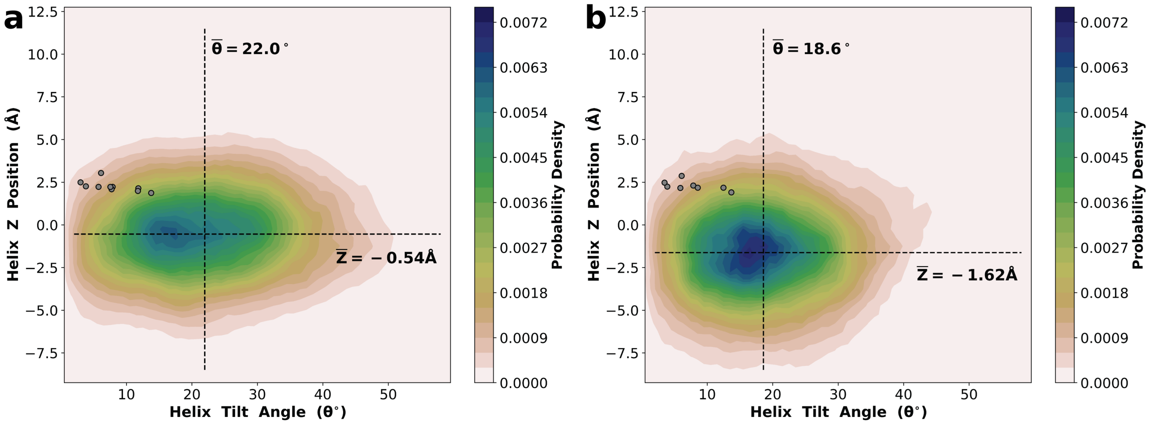


**Supplementary Figure 11. Difference in transmembrane-helix tilt angle and *z* position between ChiZ1-86 and ChiZ-FL (related to Fig. 7).** Contours display the 2-dimensional histogram in tilt angle and *z* position. Initial values after CHARMM-GUI system preparation are displayed as gray circles. (a) ChiZ1-86. (b) ChiZ-FL. The helix tilt angle is defined as the angle between the membrane normal and the vector from the first turn of the transmembrane helix (center of mass of the backbone atoms of residues 67 to 71) to the last turn (residues 82 to 86). The helix *z* position is the difference between the *z* components of the centers of mass of the transmembrane helix (residues 67 to 86) and the lipid phosphate atoms.

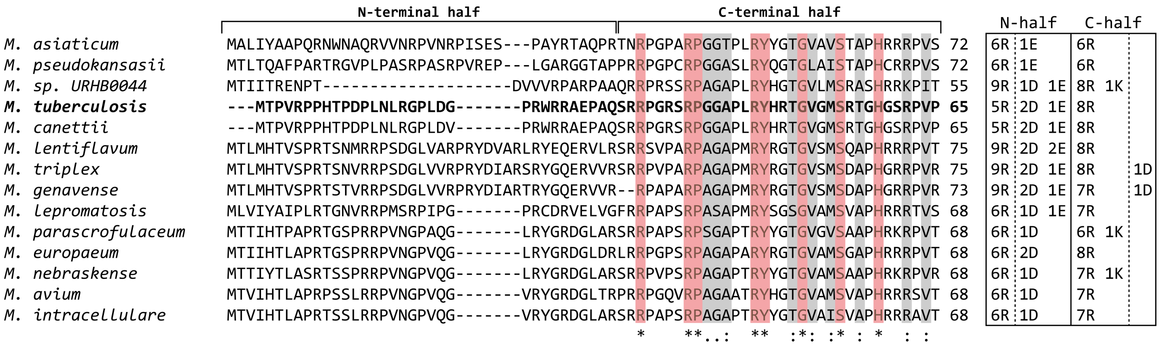


**Supplementary Figure 12. Sequence alignment and charge distribution of N-terminal regions from 14 *Mycobacteria* ChiZ homologues (related to Figs. 5-7).** Conserved and similar residues are indicated in red and grey, respectively. Counts of charged residues in N-half and C-half are shown on the right for each sequence. Multiple sequence alignment was carried out using Clustal Omega[3]. Sequences were retrieved from NCBI, with accession numbers from top to bottom: WP_036354951.1, WP_036403950.1, WP_029112638.1, WP_003899446.1, WP_015290892.1, CQD11460.1, WP_036468243.1, WP_025735491.1, WP_045842838.1, WP_040623887.1, CQD21694.1, WP_046184813.1, WP_003875224.1, and WP_014382920.1.

**Supplementary Movie 1. Dissociation and reassociation of ChiZ1-64 from one leaflet to another (related to Fig. 6).** This movie displays the portion from 20 ns to 125 ns of one 1.9-μs molecular trajectory in which ChiZ1-64 dissociated from and reassociated with membranes; snapshots at 500 ps intervals are played, at a speed of 12 snapshots per second. ChiZ1-64 is shown with N-half (residues 1-32) in yellow green and C-half (residues 33-64) in purple. Arginine side chains are shown in blue when any of its heavy atom comes to within 3.5 Å of any heavy atom in in lipids. POPG and POPE headgroups are colored in red and gray, respectively, with atom representations drawn when coming in contact with Arg residues.

**Supplementary Movie 2. Extreme fuzzy association of ChiZ1-64 with membrane (related to Fig. 5).** This movie displays the portion from 1230 ns to 1730 ns of a ChiZ1-64 trajectory reported in Fig. 5a (showing membrane contact status of individual residues and Fig. 5c (showing the snapshot at 1560 ns). Snapshots at 500 ps intervals are played, at a speed of 24 snapshots per second. $3_{10}$ helices are shown in yellow. Arginine side chains are shown in blue when any of its heavy atom comes to within 3.5 Å of any heavy atom in in lipids. POPG and POPE headgroups are colored in red and gray, respectively, with atom representations drawn when coming in contact with Arg residues.

**Supplementary Movie 3. Extreme fuzzy association of the N-terminal region of ChiZ-FL with membrane (related to Fig. 7).** This movie displays a portion from 790 ns to 1290 ns of a ChiZ-FL trajectory reported in Fig. 7d (showing the snapshot at 1161 ns). Snapshots at 500 ps intervals are played, at a speed of 24 snapshots per second. α-helices, $3_{10}$ helices, and β-sheets are shown in green, yellow, and purple, respectively. Arginine side chains are shown in blue when any of its heavy atom comes to within 3.5 Å of any heavy atom in in lipids. POPG and POPE headgroups are colored in red and gray, respectively, with atom representations drawn when coming in contact with Arg residues.
